# The CNS in the face of ART contains T cell origin HIV which can lead to drug resistance

**DOI:** 10.1101/588426

**Authors:** Gila Lustig, Sandile Cele, Farina Karim, Yashica Ganga, Khadija Khan, Bernadett Gosnell, Yunus Moosa, Rohen Harrichandparsad, Suzaan Marais, Ravindra K. Gupta, Anne Derache, Jennifer Giandhari, Tulio de Oliveira, Katya Govender, John Adamson, Vinod Patel, Alex Sigal

## Abstract

HIV persists despite antiretroviral therapy (ART) in cellular reservoirs thought to occur in distinct anatomical compartments. Therapy failure may occur because of incomplete ART adherence and possibly viral replication at some reservoir sites. The CNS may serve as a reservoir site due to lowered ART penetration and virus production from long-lived tissue resident macrophages. Compelling evidence for the CNS as a reservoir is the existence of individuals where HIV is suppressed below limit of detection in blood but detectable in the cerebrospinal fluid (CSF), termed CSF Escape. Here, we asked whether HIV in CSF Escape individuals is derived from macrophages or persists due to lowered ART. We used cell surface markers on the HIV envelope to determine the cellular source of HIV. We verified detection using *in vitro* derived virus from infected macrophages and T cells and tested CSF from CSF Escape individuals. We observed host surface markers consistent with T cell origin. We also measured ART concentrations in the CSF and plasma. We found a dramatic decrease in CSF ART concentrations described previously, but no significant difference between CSF Escape versus fully suppressed individuals. To examine the effect of the observed CSF ART concentrations on HIV replication, we used long-term infection with ART in cell culture. CSF Escape ART levels led to either HIV suppression or evolution of drug resistance, but not replication of drug sensitive HIV. These observations argue that persistent CNS viremia despite ART can be T cell generated and may result in drug resistance and therapy failure.

## Introduction

HIV persistence in the face of ART necessitates lifelong adherence to treatment. The CNS may serve as one reservoir for HIV persistence [3, 41]. HIV infection in the CNS in the absence of suppressive ART may lead to HIV-associated neurocognitive disorders (HAND). Yet, even in the presence of ART mediated suppression, there is evidence that sub-clinical cognitive impairment is common [9, 16, 18, 27, 34, 39, 40, 43, 59–61, 66–68, 75]. Another observation which supports HIV mediated effects in the face of ART in the CNS is widespread immune activation and inflammation in this compartment in the presence of therapy [1, 19, 36, 44, 79].

The elimination of the HIV reservoir in the brain may improve the quality of life of infected individuals. It may also be part of a strategy to cure HIV infection by targeting HIV reservoirs, which may be necessary across all compartments where such reservoirs persist [15, 20]. One line of evidence that the brain harbours appreciable virus in the face of ART comes from individuals who have detectable viremia in the CSF, where the CSF is used as a proxy for HIV infection in the CNS [11, 23, 49, 55], while being successfully suppressed below the level of detection in the blood.

In most cases, long-term adherence to ART does not lead to the evolution of drug resistant virus [15, 20]. One mechanism consistent with long-term persistence of ART sensitive HIV is a reservoir of latently infected CD4+ resting T cells [13–15, 20, 28, 64]. This reservoir is expected to be replenished by cellular homeostatic proliferation [12], not viral replication with the associated evolution of drug resistance [52]. This type of persistence mechanism may also involve other long-lived cell types. Examples include macrophage-linage cells such as microglia and perivascular macrophages [3, 35, 60, 61, 78] in the CNS. Tissue resident macrophages may persist for years [33, 56], and the cytotoxic effects of HIV infection on macrophages are observed to be low [4]. This would enable such infected cells to contribute to the HIV virus pool without requiring reinfection in the presence of ART. Also, cell-to-cell HIV spread originating from macrophages has been reported to have decreased ART sensitivity [22].

In addition to long-lived infected macrophage lineage cells, another reason the CNS may serve as a reservoir for HIV in the face of ART is the low drug penetration into this compartment. Reported fold-decrease in ART levels for the common reverse transcriptase inhibitor based regimen of efavirenz (EFV), emtricitabine (FTC), and and tenofovir (TFV) are approximately 200-fold, 2-fold, and 20-fold, respectively [5, 7, 8, 25]. Sufficiently decreased ARV concentrations may allow ongoing replication to occur. Whether HIV replication would lead to evolution of drug resistance would depend on sufficient selective pressure to favour drug resistant over drug sensitive virus [58]. A report describing evolution in the lymph node compartment in the face of ART [50] postulated that at sufficiently decreased drug concentrations [30], an anatomical site can serve as a sanctuary for drug sensitive virus, which allows such virus to replicate but not evolve drug resistance because ART levels are too low. Drug sensitive virus would therefore be unable to escape the sanctuary site until ART is interrupted. Such a scenario would be an explanation for the lack of evolution of drug resistance with HIV replication in the face of ART.

Here we aimed to determine if viremia detected in the CSF of CSF Escape individuals was of macrophage origin, or able to replicate as drug sensitive virus in the CNS due to lowered ART concentrations. We interrogated the CSF and matched peripheral blood of HIV infected individuals on a regimen of EFV, FTC, and TFV. We determined the cellular source of HIV by taking advantage of the fact that upon budding, HIV retains the plasma membrane of the host cell as its envelope. This contains host surface markers which may be used to detect the cellular source of the virus [46–48, 51, 74]. Our results were consistent with a T cell source of HIV in CSF Escape.

We also measured ART concentrations in the plasma and CSF using liquid chromatography–tandem mass spectrometry (LC-MS/MS) and performed evolution at the measured drug concentrations to determine if drug sensitive virus could persist. The decline in viral replication at the measured drug concentrations for individuals with CSF Escape led to a disappearance of infected cells *in vitro* over time, indicating that despite the precipitous drop in drug levels, they are nevertheless sufficiently potent to suppress HIV. Introducing an adherence gap and sequential transfer of virus from lower to higher concentrations led to the evolution of high level drug resistance. Hence, we conclude that the CNS is not a sanctuary site in the face of ART where long-term persistence of drug sensitive is possible, but rather a site where drug resistance is more likely to evolve given gaps in adherence or other factors.

## Materials and Methods

### Ethical statement

CSF and matched blood were obtained from participants indicated for lumbar puncture recruited at Inkosi Albert Luthuli Central Hospital and King Edward VIII Hospital in Durban, South Africa after written informed consent (University of KwaZulu-Natal Institutional Review Board approval BE385/13). Blood for PBMC, CD4+ and CD14+ cell isolation was obtained from adult healthy volunteers after written informed consent (University of KwaZulu-Natal Institutional Review Board approval BE022/13 and BE083/18).

### Antiretrovirals, viruses and cells

The following reagents were obtained through the AIDS Research and Reference Reagent Program, National Institute of Allergy and Infectious Diseases, National Institutes of Health: the antiretrovirals EFV, FTC, and TFV and the macrophage tropic pNL4-3(AD8) plasmid. NL4-3 and NL4-3(AD8) HIV viral stocks were produced by transfection of HEK293 cells with the molecular clone plasmids using TransIT-LT1 (Mirus) transfection reagent. Supernatant containing released virus was harvested two days post-transfection and filtered through a 0.45 micron filter(GVS) and stored in 0.5ml aliqouts at −80°C. The number of HIV RNA genomes in viral stocks was determined using the RealTime HIV-1 viral load test (Abbott, Molecular Diagnostic Services, MDS). RevCEM-E7 cells were generated as previously described [10]. Cell culture medium was complete RPMI 1640 supplemented with L-Glutamine, sodium pyruvate, HEPES, non-essential amino acids (Lonza), and 10% heat-inactivated FBS (Hyclone). PBMCs were isolated by density gradient centrifugation using Histopaque 1077 (Sigma). CD4+ or CD14+ cells were positively selected using either CD4 or CD14 Microbeads loaded onto MACS separation columns according to manufacturer’s instructions (Miltenyi Biotec). CD4+ PBMCs were grown in the above cell media supplemented with 5ng/ml IL-2 (Peprotech) and 10*μ*g/ml PHA (Sigma-Aldrich). Monocyte-derived macrophages were grown in RPMI 1640 supplemented with 10% human serum (Sigma) with added L-Glutamine, sodium pyruvate, HEPES, and non-essential amino acids (Lonza), and differentiated with 20ng/ml M-CSF (Peprotech) for 10 days.

### Generation of CD4+ PBMC and Macrophage origin virus stocks

To generate PBMC origin stocks, CD4+ PBMCs isolated and activated as described above were infected with 2*x*10^6^ RNA copies/ml NL4-3(AD8) for 24 hours. Cells were then washed 4 times in growth medium to remove the input viral stock and incubated for 4 days (approximately two full virus cycles). Virus containing supernatant was collected, centrifuged at 300g for 5 minutes to remove cells and then filtered through a 0.45 micron syringe filter (GVS) to remove cellular debris. Virus stocks were frozen as 1ml aliquots and the number of virus genomes in viral stocks was determined using the RealTime HIV-1 viral load test (Abbott). To generate monocyte-derived macrophage virus stocks, CD14+ monocytes were isolated and differentiated as described above. Cells were then infected with with 2*x*10^7^ RNA copies/ml NL4-3(AD8) for 24 hours. Cells were washed 6 times with RPMI to remove input virus, and growth media was replaced. Half the volume of media was replaced every 3 days for 10 days. Virus containing supernatant was collected, centrifuged and filtered, and viral load determined as for PBMC virus.

### Host cell marker detection on virion surface (HMV)

The following protocol was adapted from the *μ*MACS Streptavidin Kit protocol (Miltenyi): 1*μ*g of biotinylated antibodies to CD26 or CD36 (Ancell) were added to 1ml of virus, mixed and incubated for 30 minutes at room temperature. Next, 30*μ*l of strepavidin MicroBeads (Miltenyi) were added per sample, mixed and incubated at room temperature for 10 minutes. The samples were then loaded onto a *μ*Column, washed three times and bound virus eluted. Clinical virus samples were centrifuged for 13,000g for 30 seconds to clear debris before addition to antibodies. To avoid overloading columns, *in vitro* generated virus stocks from either CD4+ PBMCs or macrophages were diluted to approximately 10^4^ RNA copies/ml in PBS before addition to antibodies. The number of virus genomes in elutions from *μ*Columns was determined using the RealTime HIV-1 viral load test (Abbott, MDS).

### Surface staining for CD26 and CD36 markers and flow cytometry

Macrophages and CD4+ PBMCs were generated as described above. CD4+ PBMCs were washed once in PBS−/−. Monocyte-derived macrophages were washed once in PBS−/− then incubated in 5mM EDTA in PBS−/− for 30 minutes on ice. Macrophages were collected by pipetting vigorously and the remaining attached cells were removed by gentle scraping. Cells were then incubated with CD3-APC or CD68-APC and one of CD26-FITC, CD26-PE, CD36-FITC and CD36-PE (Biolegend) in staining buffer (PBS−/− with 3%FCS) for 30 minutes on ice. The samples were then washed, resuspended in 400*μ*l staining buffer and acquired on a FACSCalibur machine (BD Biosciences). Results were analyzed with FlowJo software.

### Generation of YFP-NL4-3(AD8)

pNL4-3(AD8) was used as the source of the macrophage tropic R5 AD8 HIV envelope which was cloned into the pNL4-3-YFP vector (gift from David Levy), replacing the NL4-3 X4 specific Env. between the unique EcoRI-BamHI restriction sites. The fragment of ENV was excised from pNL4-3(AD8) using BamHI and EcoRI restriction enzymes (NEB) and ligated using T4 ligase (Invitrogen) into the EcoRI-BamHI restricted pNL4-3-YFP backbone to produce YFP-NL4-3(AD8).

### Detection of ART concentrations in CSF and matched plasma by LC-MS/MS

Sample analysis was performed using an Agilent High Pressure Liquid Chromatography (HPLC) system coupled to the AB Sciex 5500, triple quadrupole mass spectrometer equipped with an electrospray ionization (ESI) TurboIonSpray source. The LC-MS/MS method was developed and optimised for the quantitation of drug analytes; tenofovir, emtricitabine, efavirenz, lopinavir, ritonavir, nevirapine, zidovudine, lamivudine, abacavir, atazanvir and dolutegravir. A protein precipitation extraction method using acetonitrile was used to process clinical plasma and CSF samples. The procedure was performed using 50 *μ*l of plasma or CSF sample, 50 *μ*l of water and 50 *μ*l of ISTD solution was added and the sample was briefly mixed. 150 *μ*l of acetonitrile was subsequently added to facilitate protein precipitation, vortex mixed and centrifuged at 16000g for 10 minutes at a temperature of 4°C. 170 *μ*l of the clear supernatant was then transferred to a clean micro-centrifuge tube and dried down using a SpeedVac dryer set at 40°C. The dried samples were then reconstituted in 100 *μ*l of 0.02% SDC (sodium deoxycholate) in Millipore filtered water, vortex mixed, briefly centrifuged, placed in a small insert vial, capped, placed in the auto sampler compartment (maintained at 4°C) and analysed using LC-MS/MS. The analytes were separated on an Agilent Zorbax Eclipse Plus C18 (2.1 × 50mm, 3.5 *μ*m) HPLC column using gradient elution. The column oven was set at 40°C, a sample volume of 2 *μ*l was injected and the flow rate was set to 0.2 ml/min. Mobile phase A consisted of water with 0.1% formic acid and B consisted of acetonitrile with 0.1% formic acid. The drug analytes were monitored using multiple-reaction monitoring mode for positive ions except for efavirenz which was monitored in the negative ion scan mode. Analyst software, version 1.6.2 was used for quantitative data analysis. The LC/MS/MS method showed linearity and allowed for accurate quantitation across a concentration range of 180 – 6000 ng/ml for lopinavir, ritonavir, nevirapine, abacavir, lamivudine, zidovudine, atazanavir and dolutegravir, 150 – 5000 ng/ml for efavirenz, 90 – 3000 ng/ml for tenofovir and 24 – 780 ng/ml for emtricitabine.

### Calculation of expected drug concentrations after 72 hour adherence gap (EFV tail)

To calculate the expected drug concentrations after a 72 hour adherence gap, we used the previously reported half-lives (*t*_1/2_) for EFV, FTC, and TFV [2, 77] of 48 hours, 10 hours, and 14 hours respectively. The decay rate in hours (*r*) was calculated per drug as *r* = *ln*2/*t*_1/2_ and the remaining drug after 72 hours calculated as *D*_72*h*_ = *D*_0_*e*^−*rt*^, where *t* = 72 hours, and *D*_0_ is the median CSF concentration of the drug in CSF Escape individuals.

### *in vitro* evolution

RevCEM-E7 reporter cells grown at a concentration of 10^6^ cells/ml in 4ml of growth medium in 6-well plates (Corning) were infected with 2*x*10^7^ RNA copies NL4-3 HIV per ml. Cells were passaged in the presence of 3 concentrations of EFV, FTC, and TFV using a dilution of 1:2 every 2 days, where half the cell culture was removed and fresh medium with drugs was added.The three drug concentrations used were: 1) EFV tail, 18 ng/ml EFV; 2) CSF median, 51 ng/ml EFV, 190ng/ml FTC, 7 ng/ml TFV; and 3) plasma median, 2200 ng/ml EFV, 194 ng/ml FTC and 66 ng/ml TFV. Proliferation of uninfected cells was sufficient to maintain uninfected target cell numbers. The removed fraction of cells was used to detect infection by flow cytometry using a timed acquisition on a FACSCaliber machine. The concentration of infected cells was multiplied by the volume and cumulative dilution factor to obtain the number of infected cells per well.

### Illumina Sequencing

Genomic extraction was preformed on 10^5^ – 10^6^ cells from the *in vitro* evolution experiments using the *Quick*-DNA Miniprep Plus Kit (Zymo Research). The gDNA was then diluted to a range between 100-200ng/5*μ*l in TE buffer. Amplification of a 500bp region of the reverse transcriptase was done with 0.25*μ*l ExTaq (TaKaRa), 5*μ*l of 10x Ex Taq buffer, 4*μ*l 2.5mM of each dNTP, 5*μ*l gDNA temple and 1*μ*M of each primer with the final reaction volume 50*μ*l. Cycling program was 30 cycles of denaturing at 98°C for 10 seconds, annealing at 55°C for 30 seconds and elongation at 72°C for 1 minute. The forward primer sequence was 5’-tcgtcggcagcgtcagatgtgtataagagacagTTAATAAGAGAACTCAAGATTTC-3’ and reverse primer sequence 5’-gtctcgtgggctcggagatgtgtataagagacagCAGCACTATAGGCTGTACTGTC-3’, where capitalized end regions are adaptors for Illumina sequencing. Products were visualized on a 1% agarose gel and bands correponding to 500bp were excised with an x-tracta Gel Extraction Tool (Sigma). The DNA was extracted from the gel using the Gel Extraction Kit (Qiagen). Illumina sequencing was then performed. Input DNA was quantified using the Qubit dsDNA HS Assay system and diluted in molecular-grade water to reach the starting concentration of 0.2 ng//*mu*l. Barcodes were added to each sample using the Nextera XT Index kit (Illumina, Whitehead Scientific, SA). Barcoded amplicons were purified using Ampure XP beads (Beckman Coulter, Atlanta, Georgia) and fragment analysis was performed using the LabChip GX Touch (Perkin Elmer, Waltham, US). The library was pooled at a final concentration of 4nM and further diluted to 3pM. The library was spiked with 20% PhiX plasmid which served as an internal control to account for low diversity libraries. The spiked library was run on a Miseq v2 with a Nano Reagent kit.

## Results

### HIV in CSF of CSF Escape individuals is of T cell origin

We have sampled CSF and matched blood from ART treated participants (n=139) clinically indicated for lumbar puncture as part of their diagnostic workup (Table 1). 73 (53%) of this group showed an undetectable viral load (VL) in the plasma. Out of the subgroup of successfully suppressed individuals in the plasma, 16 (22%) had detectable viremia in the CSF termed CSF Escape [23].

**Table 1.** Participant details.

| State | n (F/M) | Age | VL CSF | VL Plasma | CD4 count | Co-infect |
| --- | --- | --- | --- | --- | --- | --- |
| CSF<br>Escape | 16<br>(8/8) | 39<br>(34 – 49) | 2915 ± 15<br>(977 – 21045) | < 40 | 474<br>(272 – 660) | 31% |
| Suppressed | 57<br>(37/20) | 34<br>(28 – 42) | < 40 | < 40 | 581<br>(340 – 721) | 35% |
| Viremic | 66<br>(45/21) | 34<br>(30 – 41) | 2760 ± 15<br>(< 40 – 42355) | 7923 ± 15<br>(373 – 48654) | 266<br>(103 – 964) | 39% |

For each sample, we used LC-MS/MS to assay for the presence of ART regimen components. This consisted of the South African first line regimen of EFV, FTC, and TFV as well as the protease inhibitors lapinovir and ritonavir (LPV/r) and the reverse transcriptase inhibitor nevirapine (NVP). We excluded samples with LPV/r and NVP so that only samples from individuals on first line therapy were analyzed further.

We modified an existing approach [46–48, 51, 74] to interrogate the cellular origin of cell-free HIV by detecting host cell markers on the surface of the HIV virion, an approach we term Host Marker on Virion detection (HMV, Fig 1A). As the virus buds out of the infected cell, it uses the cellular plasma membrane as its envelope [71]. The cell membrane contains host surface markers which are retained by the virion on its surface and can be bound with antibodies. We coupled the antibodies to magnetic beads and used a VL assay to determine the number of bound virions after washing off unbound virions (Fig 1A). Using this technique, we successfully detected two cell surface markers associated with cell type: CD26, a dipeptidyl-peptidase involved in T cell activation and preferentially expressed by CD4+ T cells [29, 54], and CD36, a scavenger receptor involved in recognition and phagocytosis of phospholipids and lipoproteins and expressed in macrophages as well as other cell types [57, 65]. These markers have been previously shown to be effective at differentiating between CD4+ T cell and macrophage origin HIV [51, 74].

**Figure 1.**
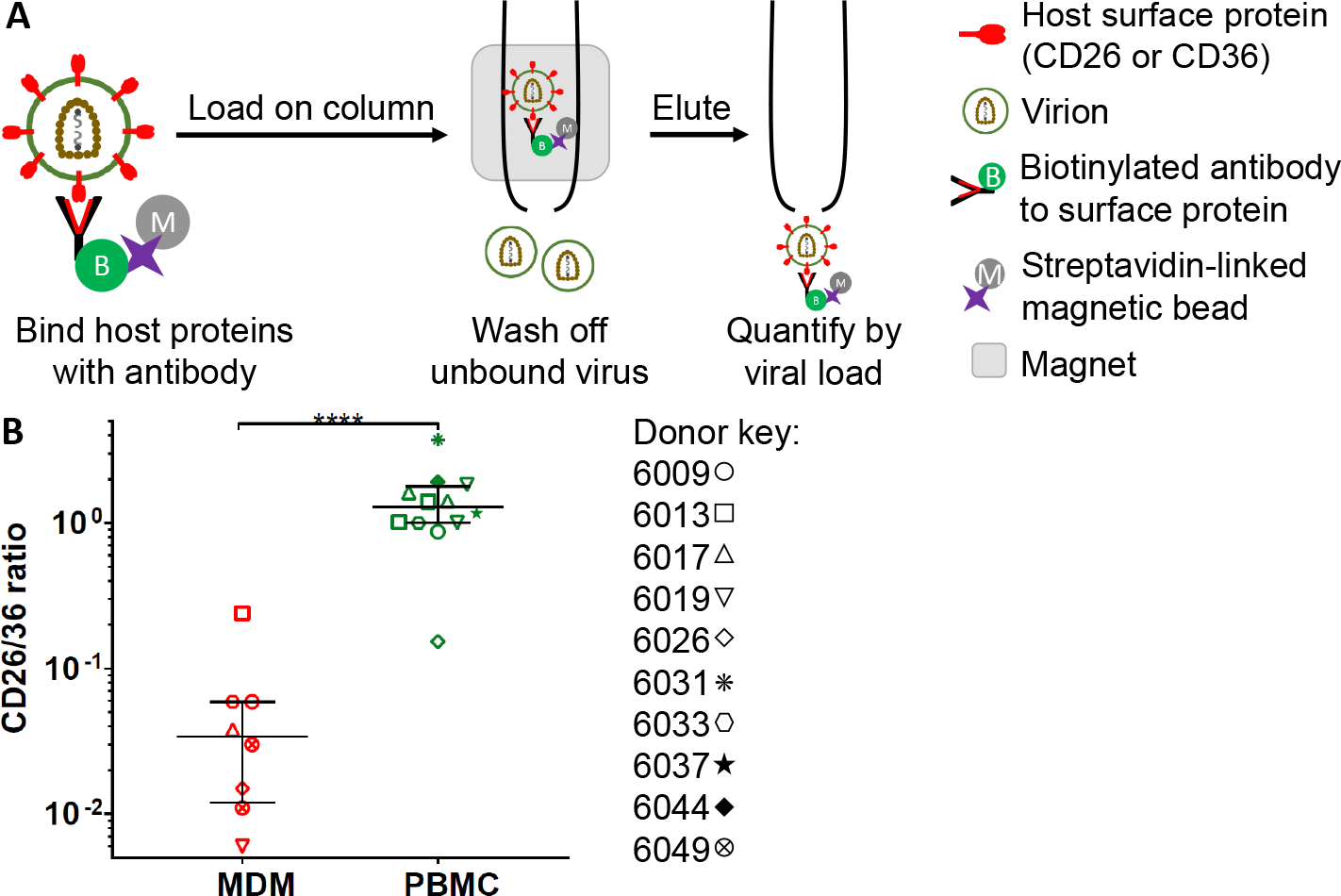
HMV detection of cellular origin of HIV. **A**, Schematic of method. Cell-free virus from cell suspensions or plasma/CSF was bound to columns with CD26 or CD36 beads. After washing off unbound virus, virus bound to the columns was eluted and quantified using a VL assay. **B**, Monocyte derived macrophages or CD4+ PBMCs were infected with NL4-3(AD8) macrophage tropic HIV and supernatant from infected cells was loaded on columns to quantify CD26/CD36 ratio. Shown are median and IQR of values derived from different blood donors (symbols on right, associated number is participant identification number). The CD26/36 ratio of MDM derived virus was significantly lower than that derived from PBMCs (p < 0.0001, Mann-Whitney non-parametric test).

We used the ratio of CD26 to CD36 to determine the virus producing cell type. To verify that HMV detection is effective at differentiating between macrophage and T cell sources of HIV, we infected either monocyte derived macrophages (MDMs) or CD4 expressing peripheral blood mononuclear cells (PBMCs) with the macrophage tropic NL4-3(AD8) HIV strain which infects both CD4 T cells and macrophages. Confirming previous results [51, 74], there was a significant increase (*p* < 0.0001) in the CD26/CD36 ratio in virus originating from CD4+ PBMCs relative to MDMs (Fig 1B), driven primarily by the increase in CD26 positive virions (SFig 1). The median increase was over one order of magnitude, from 0.034 (IQR 0.012 - 0.059) for MDMs to 1.28 (IQR - 1.78) for PBMCs. Using YFP labelled macrophage tropic HIV (YFP-NL4-3(AD8)), we observed no infection of CD14+ monocytes in PBMCs, indicating that the PBMC derived virus originated in CD4+ T cells (SFig 2).

**Figure 2.**
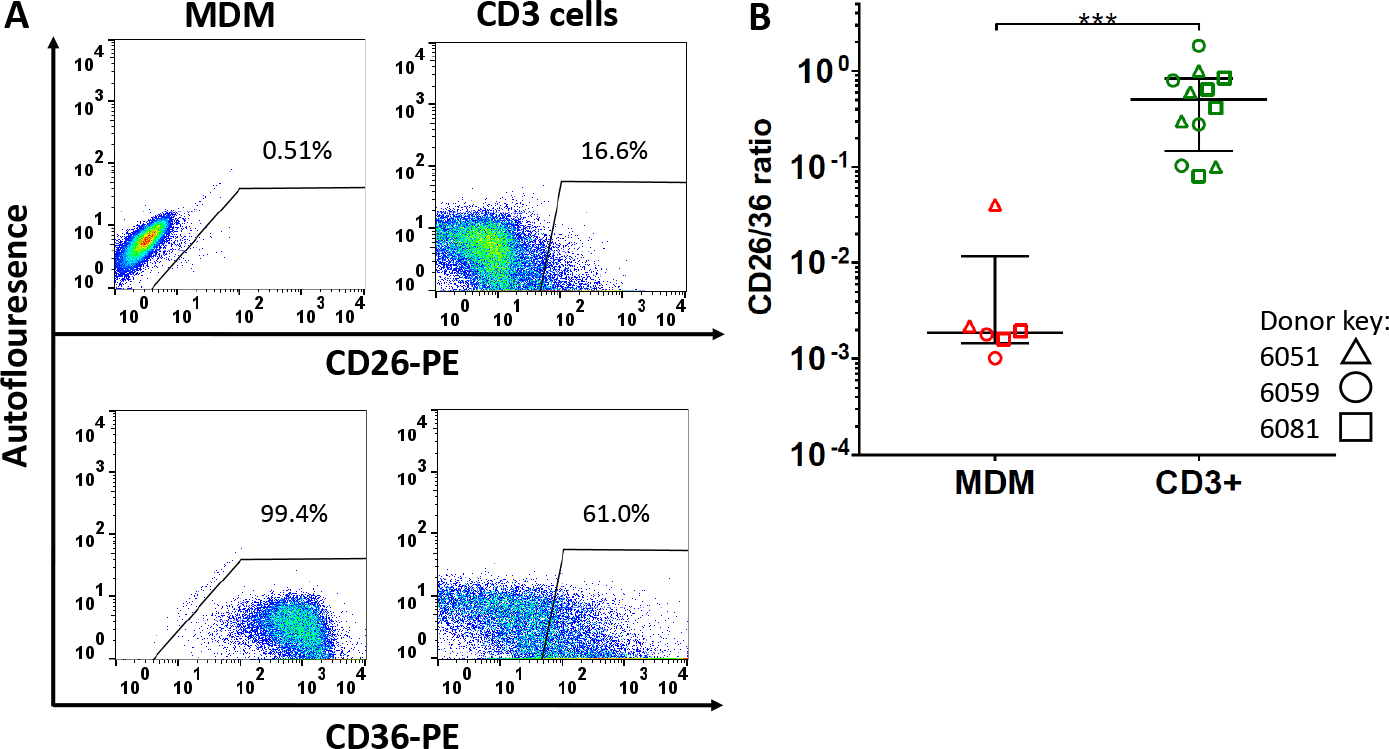
Surface expression of CD26 and CD36 on CD3+ T cells and MDM. **A**, Flow cytometry of MDMs, pre-gated on CD68 (left panels) or CD3 (right panels) stained for CD26 or CD36. **B**, Surface expression of CD26 and CD36 on MDM or CD3+ T cells from three donors (symbols on right, associated number is participant identification number). Median and IQR range shown. MDM ratio was significantly lower (p < 0.0001, Mann-Whitney non-parametric test).

We used cell surface expression to confirm that T cells expressed a higher ratio of CD26 to CD36 relative to MDMs. We detected CD26 and CD36 on the surface of MDMs and CD3+ T cells from PBMCs (Fig 2A). MDMs uniformly expressed CD36, with little CD26 expression. In contrast, both CD26 and CD36 were expressed on CD3+ T cells for CD3+ T cells (*p* < 0.0001, Fig 2B). The T cell ratio of CD26 to CD36 surface expression was 0.505 (IQR 0.83-0.15) while that of MDMs was 0.002 (IQR 0.01-0.0015). The surface expression CD26/CD36 ratio is consistent with that obtained by detecting these surface markers on virions.

We next used HMV detection to infer the cellular origin of *in vivo* derived CSF Escape virus. We measured the number of virions bound per CD26 or CD36 column from the same CSF sample and compared the CD26/CD36 ratios from CSF Escape individuals with the *in vitro* values for MDMs and CD4+ PBMCs (Fig 3). The median CSF escape CD26/CD36 ratio, at 1.2 (IQR 0.99 - 2.51), was significantly higher than the ratio obtained from MDM infection (*p* < 0.0001). It was not significantly different from PBMC T cell derived virus. We therefore conclude that the surface markers on HIV found in the CSF of individuals with CSF escape is consistent with a T cell origin.

**Figure 3.**
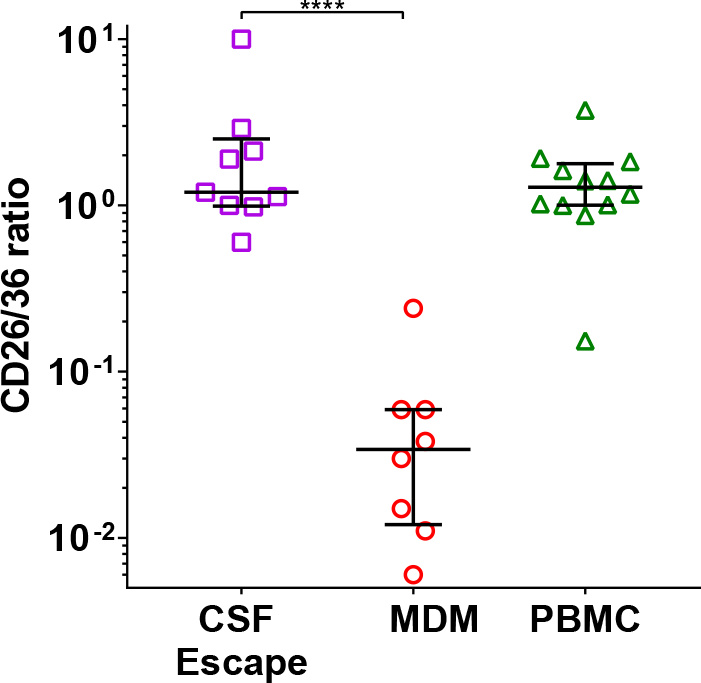
in vivo cellular source of HIV from participants with CSF Escape. CSF samples from CSF Escape individuals were split into two, with half the sample applied to a column with CD26 antibodies and the other to a column with CD36 antibodies to obtain the ratio of virus bound by each. Median ratio of CSF Escape virus (left) was compared to median ratios from in vitro derived virus from infected MDMs (middle) or PBMCs (right) and was found to be significantly higher than the MDM ratio but not different from the ratio of PBMC derived virus (p < 0.0001, p > 0.5 respectively, Mann-Whitney non-parametric test).

We also compared the virus from viremic study participants, both from the CSF and plasma, to that of participants with CSF escape (Fig 4). We obtained no significant differences between these groups, though interestingly the plasma ratios showed a distribution with a considerable number of values in the lower range relative to either the viremic or escape CSF, indicating that macrophage lineage cells may contribute to the viral pool in the plasma in a subset of viremic individuals.

**Figure 4.**
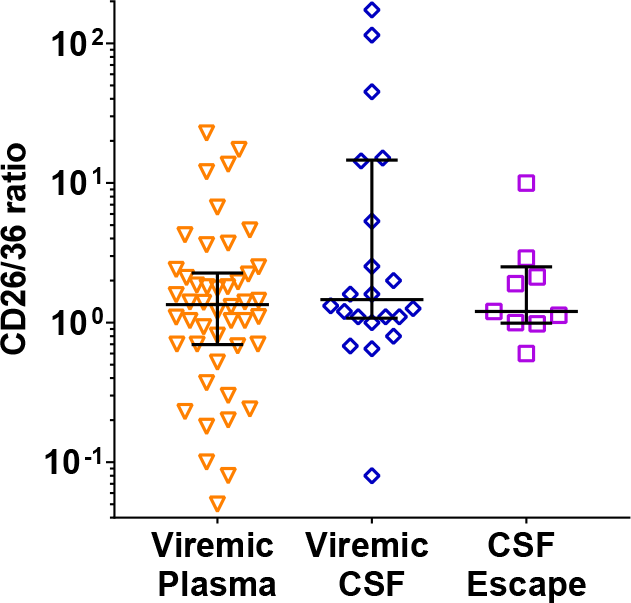
Cellular source of HIV in CSF Escape versus viremic participants. CD26 to CD36 ratios of plasma virus from viremic individuals (left) or CSF virus from viremic individuals (middle) were found to be not significantly different from CSF virus from CSF Escape (right) (Kruskal-Wallis test with Dunn multiple comparisons corrections).

### ART levels in CSF similar between individuals with CSF Escape and full suppression

To investigate if the CNS can serve as a sanctuary site which allows replication of drug sensitive HIV [50], we measured the plasma and CSF concentrations of ART components for individuals on the first line regimen of EFV, FTC and TFV. There was a precipitous drop in EFV concentrations from the plasma to the CSF in suppressed and CSF escape individuals (SFig 3), consistent with previous studies [5, 6, 72]. The drop in EFV concentrations in viremic individuals was less pronounced and the EFV concentrations in the plasma of the viremic group bifurcated into individuals with low and high levels of EFV, likely the result of a subset of individuals with low adherence and a second subset with virologic failure due to evolution of drug resistance (SFig 3). Again consistent with previous studies, there was also a drop in FTC and TFV concentrations in the CSF versus plasma, though this decrease was more attenuated for FTC. Interestingly, in participants with CSF escape, there were no significant difference between FTC concentrations in plasma versus CSF (SFig 3).

We compared drug concentrations between viremic, suppressed, and CSF Escape participants. With every drug and in every compartment, viremic participants had significantly lower drug concentrations compared to suppressed and CSF Escape individuals (Fig 5). This indicates that there is a subset of viremic participants with low adherence, and a sub-population of participants with low drug levels in the viremic group can be clearly seen in the plasma compartment for all three drugs and in the CSF compartment with FTC. However, there was no significant difference between CSF levels of EFV, FTC, and TFV between the suppressed and CSF Escape groups (Fig 5). There was also no significant difference in plasma levels of all three drugs. This indicates that it is unlikely that a reduction in CSF drug levels in CSF Escape individuals is responsible for detectable virus in the CSF compartment.

**Figure 5.**
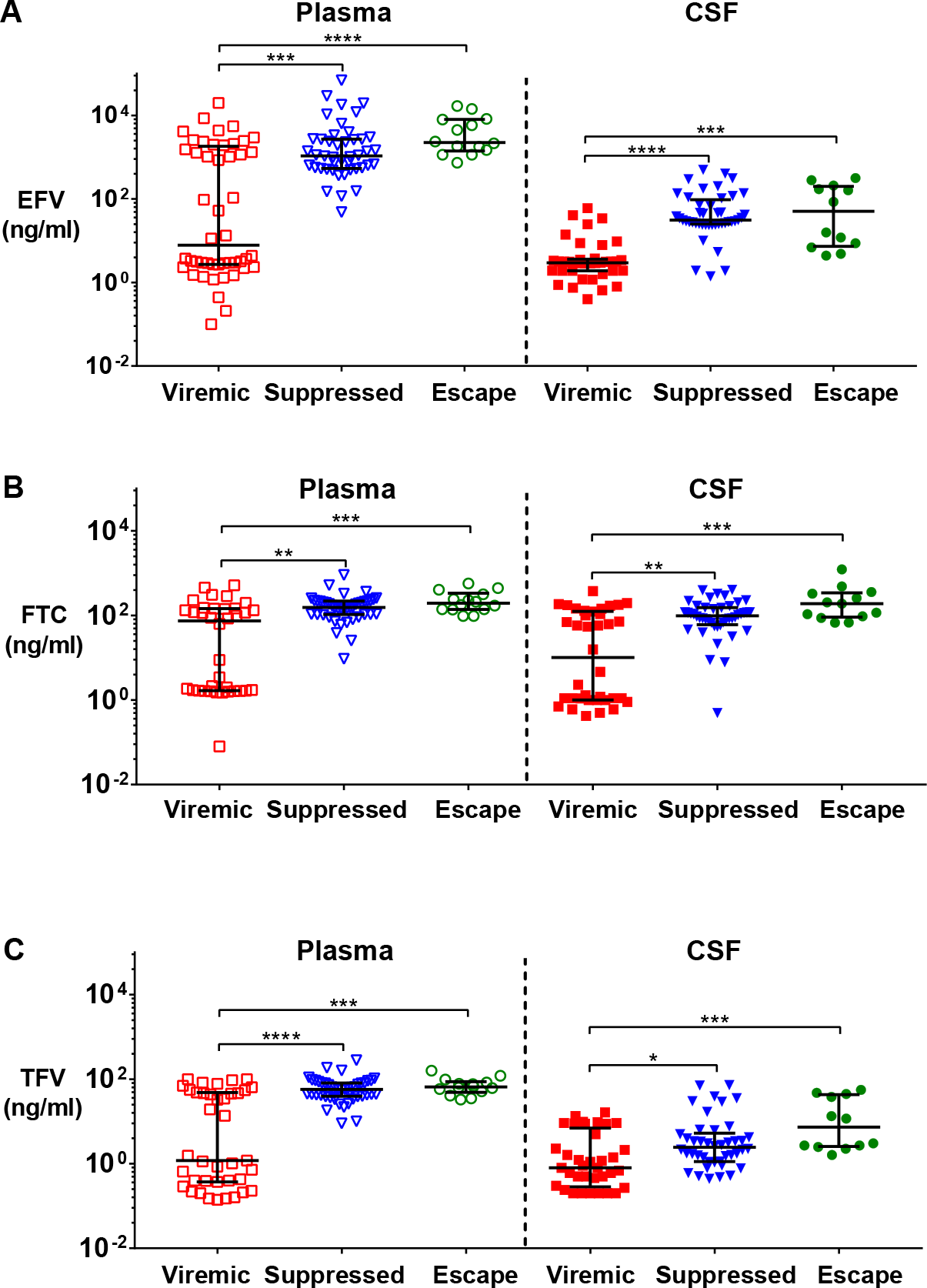
ART concentrations in plasma and CSF are similar in between suppressed and CSF Escape individuals. **A**, EFV concentrations in the plasma (left) and CSF (right) in viremic, suppressed, and CSF Escape participants. Median EFV concentration in the plasma was 7.8 ng/ml (IQR 2.7-1833 ng/ml) in viremic, 1080 ng/ml (IQR 538-2720 ng/ml) in suppressed, and 2250 ng/ml (IQR 1403-8048 ng/ml) in CSF Escape. Median EFV concentration in the CSF was 2.9 ng/ml (IQR 1.9-3.6 ng/ml) for viremic, 31.1 ng/ml (IQR 25.2-96.3 ng/ml) for suppressed, and 50.7 (IQR 7.3-201 ng/ml) for CSF Escape. The concentrations in suppressed and CSF Escape were significantly higher both in the plasma and CSF versus the viremic group (p = 0.0009 and p < 0.0001 respectively for plasma and p < 0.0001 and p = 0.0001 for CSF). However, there was no significant difference between suppressed and CSF Escape EFV concentrations in either the plasma of CSF (p = 0.26 and p > 0.99 respectively). **B**, FTC concentrations in the plasma (left) and CSF (right) in viremic, suppressed, and CSF Escape participants. Median FTC concentration in the plasma was 74.1 ng/ml (IQR 1.66-144 ng/ml) in viremic, 155 ng/ml (IQR 105-218 ng/ml) in suppressed, and 194 ng/ml, (IQR 139-337 ng/ml) in CSF Escape. Median FTC concentration in the CSF was 10.1 ng/ml (IQR 1.0-124 ng/ml) for viremic, 98.3 ng/ml (IQR 60.6-153 ng/ml) for suppressed, and 190 ng/ml (IQR 92.1-342 ng/ml) for CSF Escape. The concentrations in suppressed and CSF Escape were significantly higher both in the plasma and CSF versus the viremic group (p = 0.001 and p = 0.0005 respectively for plasma and p = 0.005 and p = 0.0002 for CSF). There was no significant difference between suppressed and CSF Escape EFV concentrations in either the plasma of CSF (p = 0.63 and p = 0.19 respectively). **C**, TFV concentrations in the plasma (left) and CSF (right) in viremic, suppressed, and CSF Escape participants. Median TFV concentration in the plasma was 1.2 ng/ml, (IQR 0.4-48.3 ng/ml) in viremic, 57.2 ng/ml (IQR 39.2-81.2 ng/ml) in suppressed, and 65.7 ng/ml (IQR 48.7-87.2 ng/ml) in CSF Escape. Median TFV concentration in the CSF was 0.8 ng/ml (IQR 0.3-6.9 ng/ml) for viremic, 2.4 ng/ml (IQR 1.1-5.3) for suppressed, and 7.4 ng/ml (IQR 2.6-43.1) for CSF Escape. The concentrations in suppressed and CSF Escape were significantly higher both in the plasma and CSF versus the viremic group (p < 0.0001 and p = 0.0004 respectively for plasma and p = 0.02 and p = 0.0004 for CSF). There was no significant difference between suppressed and CSF Escape EFV concentrations in either the plasma of CSF (p > 0.99 and p = 0.16 respectively). Significance was measured using the Kruskal-Wallis with Dunns’ corrections for multiple comparisons for each compartment separately.

### ART levels in the CSF of individuals with CSF Escape are sufficient to exert selective pressure for evolution of drug resistance

We asked whether the observed decrease in ART levels in the CSF in individuals with CSF Escape were sufficient to enable the CSF act as a sanctuary site where drug sensitive HIV can persist [50]. We therefore passaged drug sensitive HIV in vitro at the measured CSF Escape median CSF and plasma concentrations of EFV, FTC, and TFV. When passaging was done at the plasma median concentrations of EFV (2200 ng/ml), FTC (194 ng/ml), and TFV (66 ng/ml), it led to a decline in infected cell numbers. Infected cells were below the dilution corrected level of detection after 22 ± 3 days in culture (Fig 6A). The same experiment was performed at the CSF median drug levels of EFV (51 ng/ml), FTC (190 ng/ml), and TFV (7 ng/ml). Similarly to plasma drug concentrations, the number of infected cells declined until it was undetectable at 26 ± 3 days in culture (Fig 6B). In agreement with the predicted effects of the drugs on viral replication [62], this indicates that ART levels in the CSF of CSF Escape individuals are sufficient to result in a decay in infected cell numbers.

**Figure 6.**
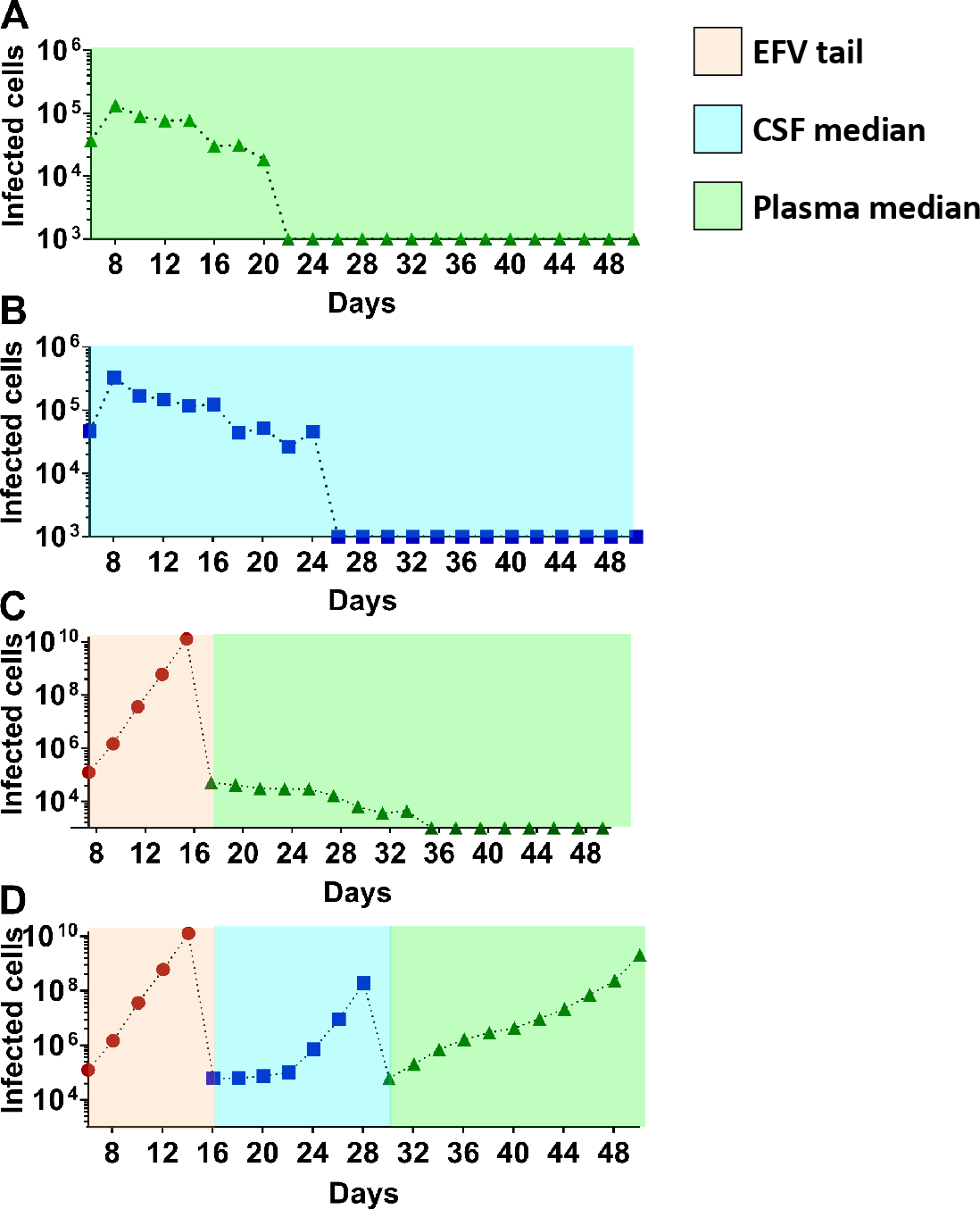
Decline in infected cell numbers at ART levels found in plasma and CSF of participants with CSF Escape. Dilution corrected number of infected cells through time in the face of: **A**, Median plasma ART concentrations. **B**, Median CSF ART. **C**, Calculated ART concentration derived after a 72-hour adherence gap (EFV tail), then transfer of infected cells to the median plasma ART concentration. **D**, Replication at the EFV tail ART concentration, then transfer to CSF median ART concentration, then further transfer to the plasma median ART concentration.

We next calculated the estimated ART levels (Materials and Methods) after a 3 day adherence gap. Interruptions on this timescale occur in approximately 20% of individuals on ART monthly [32]. Due to the longer half-life of EFV relative to FTC and TFV [2, 77], such an interruption effectively leads to EFV monotherapy [73]. We refer to this drug condition as “EFV tail”. The EFV tail drug condition led to viral replication and the evolution of the L100I EFV resistance mutant (Fig 6C-D, Fig 7). Transfer of cells infected with the evolved virus to the median plasma ART concentration led to a decay in the number of infected cells (Fig 6C). In contrast, transfer to the median CSF ART concentrations led to an increase in infected cell numbers (Fig 6D) and the evolution of FTC resistance and further resistance to EFV (Fig 7). Transfer of this virus to the plasma ART concentration led to an increase in infected cell numbers (Fig 6D) and high level resistance. The mutations which arose at these stages were the M184V or M184I FTC resistance mutations during median CSF Escape ART levels, and the K103N high level EFV resistance mutation either at median CSF Escape ART levels (2 experiments) or plasma median ART (1 experiment). In the experiments where the K103N mutation arose early, it temporarily or permanently supplanted the L100I mutation (Fig 7B-C).

**Figure 7.**
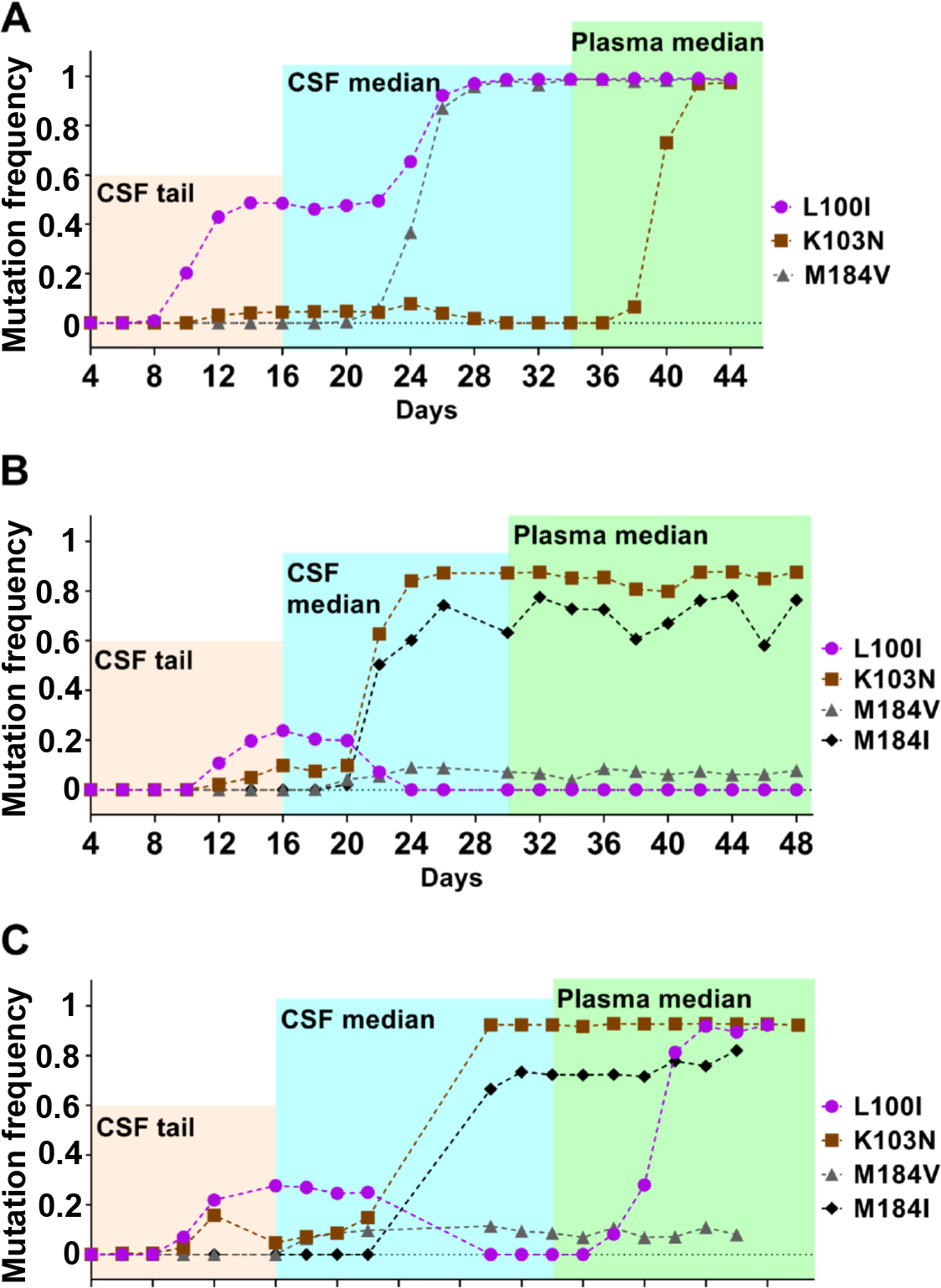
Evolution of drug resistance with sequential increase in ART levels. Accumulation of mutations as detected by Illumna sequencing for three independent experiments **(A-C)**, with infection levels in experiment **(A)** shown in Figure 6D. L100I is a low level resistance mutation to EFV, M184V and M184I are high level resistance mutations to FTC and K103N is a high level resistance mutation to EFV according to the HIV Drug Resistance Database (https://hivdb.stanford.edu/).

These results predict that ART levels measured in the CSF of CSF Escape individuals will lead to decay in infected cell numbers or, in the case of a gap in adherence, the evolution of drug resistant HIV. This prediction is supported by several studies showing drug resistance mutations in the face of ART in CSF from CSF Escape and viremic individuals [69, 76] and our own limited sequencing (STable 1). Here, one of the CSF Escape HIV sequences was drug sensitive. However, the CSF sample was collected close to the ART start date, suggestive of a previously reported longer decay time of drug sensitive HIV in the CSF compartment [24].

## Discussion

The presence of individuals with CNS Escape, where HIV is fully suppressed peripherally but present in detectable levels in the CSF, suggests the possibility that, at least in some individuals, the CNS can serve as a an HIV reservoir site either because of HIV production from long lived tissue resident macrophages, or because of decreased ART penetration leading to ongoing replication. Here, we tried to establish whether macrophages were indeed the source of viremia in study participants with CSF Escape, or whether drug levels in the CSF were low enough in CNS Escape for ART sensitive HIV to replicate.

We used the ratio of the cell surface markers CD26 and CD36 to differentiate between virus of T cell versus macrophage origin. This technique is based on the much lower expression of CD26 on the cell surface of macrophages combined with a higher expression of CD36 on these cells relative to T cells. It was effective at discriminating between T cell and macrophage origin virus *in vitro*. The results obtained from CSF HIV from CSF Escape participants were consistent with a T cell origin, decreasing the likelihood that infected macrophages are the drivers of HIV production in the CSF in this group of individuals.

We confirmed the previously reported lower ART penetration into the CSF compartment in our study population, which consisted of predominantly HIV Clade C infected participants [37]. However, the drop in drug concentrations in the CSF relative to the peripheral blood was similar between individuals with CSF Escape and those who were fully suppressed in both compartments, ruling out lower ART concentrations in the CSF as the reason for CSF Escape. While drug concentrations were lowered in the CSF, they were sufficient to suppress replication by drug sensitive HIV *in vitro*.

Only an ART concentration where an adherence gap was assumed led to an increase in infected cell numbers due to evolution of drug resistance *in vitro*. In this case, replication at CSF ART concentrations was a necessary step to evolve sufficient resistance for the virus to transition to expanding infection and further evolution of drug resistance mutations at plasma ART concentrations.

These results are consistent with a role for the CNS in the face of ART as a transient reservoir where virus either decays or evolves drug resistance and possibly results in virologic failure. There are several caveats to these conclusions. First, virus from *in vivo* infected microgila, the tissue resident macrophages in the brain thought to be an infection target of HIV [31, 38, 45], may show different surface marker profiles relative to the monocyte-derived macrophages we used as a standard. While this is a possibility, much is known of the function of the CD36 scavenger receptor in microglia [17, 26, 42, 53, 70], while a rare characterization of CD26 in these cells reports them to be negative [21]. Second, we only tested the CSF, while HIV can persist in additional CNS compartments [3]. This is a limitation of the current study. However, virus which is strongly compartmentalized and unable to enter the CSF may have difficulty exiting the CNS and seeding infection in the periphery on the two week timescale which is usual for rebound of infection after ART interruption [15, 63]. Hence, it may not be a reservoir which is the source of HIV rebound.

Based on the data presented here, we propose that the CSF may be a transient reservoir which tracks adherence gaps and possibly other factors which permit viral replication in the CNS. Replication in this compartment would then decay or lead to virologic failure due to evolution of drug resistance. This shows that HIV reservoirs may be complex, with different reservoir sites supporting different mechanisms of persistence.

## Acknowledgments

This work was supported by National Institutes of Health grant R21MH104220 to AS.

---

**SFigure 1.**
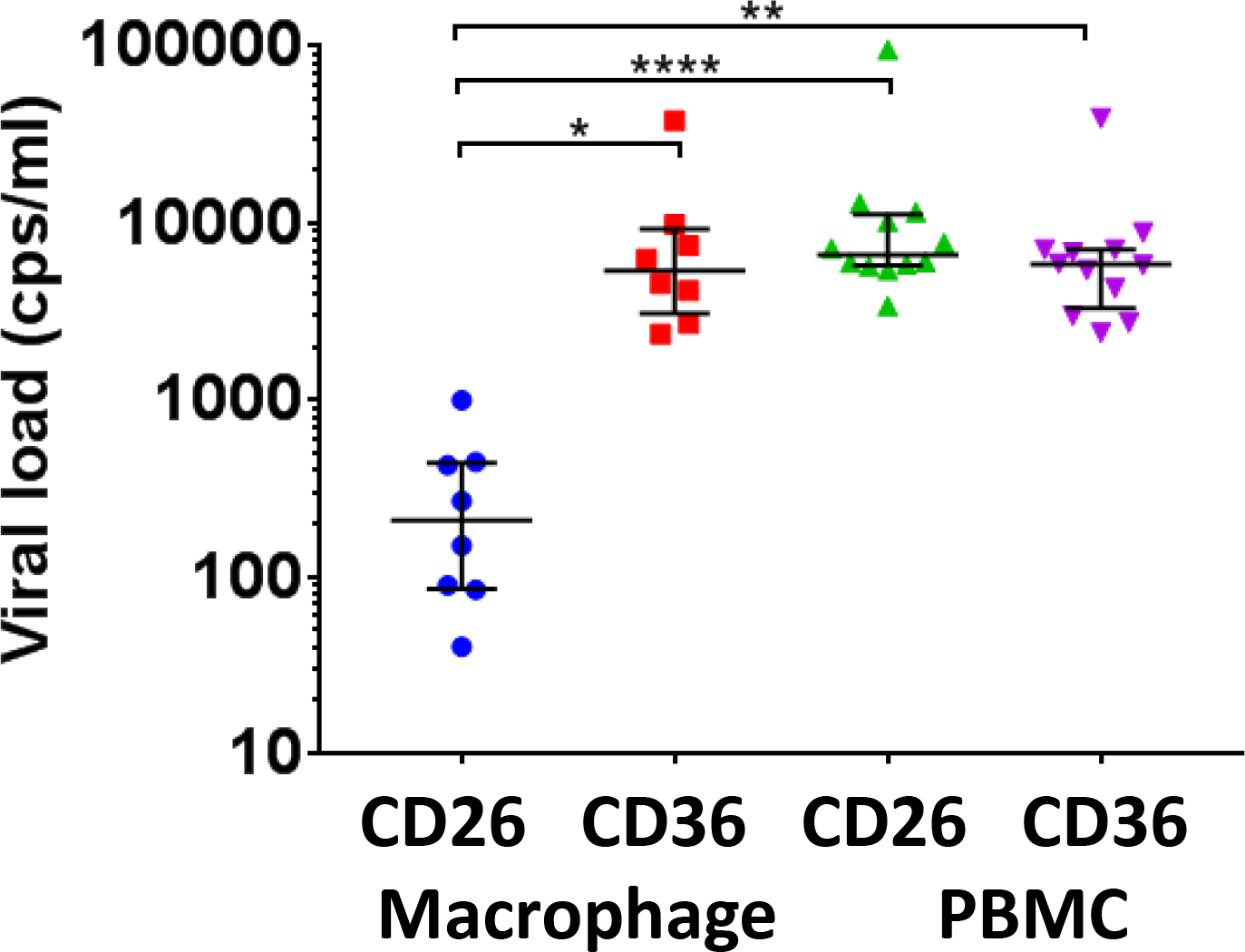
Absolute RNA copies from CD26 versus CD36 columns in *in vitro* infection. MDMs or CD4+ PBMCs were infected with NL4-3(AD8) macrophage tropic HIV and 1ml supernatant diluted to approximately 10^4^ HIV RNA copies/ml from infected cells was loaded on columns to quantify CD26 and CD36. Shown are median and IQR for different blood donors. MDM values: CD26 median 208 HIV RNA copies/ml (IQR 85-440 copies/ml), CD36 median 5403 copies/ml (IQR 3076-9263 copies/ml). PBMC values: CD26 median 6600 copies/ml (IQR 5749-11185 copies/ml), CD36 median 5862 copies/ml (IQR 3309-5862 copies/ml). Median level of MDM CD26 is significantly lower from than MDM median CD36 (*p* < 0.05), PBMC median CD26 (*p* < 0.0001), and PBMC median CD36 (*p* < 0.001). Kruskal-Wallis non-parametric test with Dunn multiple comparisons correction.

**SFigure 2.**
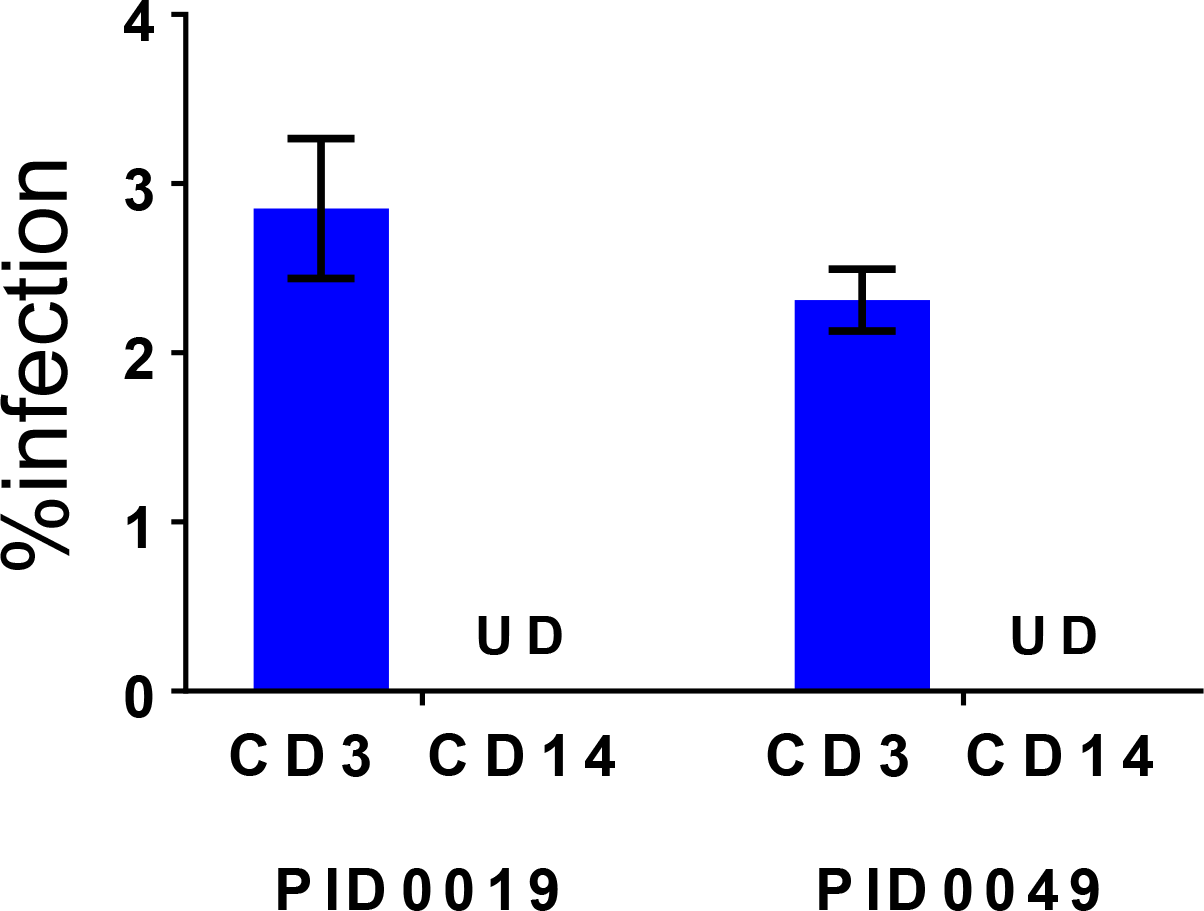
Infection of CD3+ and CD14+ cells with macrophage tropic HIV. PBMCs were infected with 2*x*10^7^ RNA copies/ml YFP-NL4-3(AD8). 2 days post-infection, cells were collected and stained with CD3 and CD14 antibodies, then analyzed for infection by detection of YFP positive cells in the CD3+ and CD14+ populations using flow cytometry. %infection shown is from the CD3 or CD14 populations. PID, participant identification number. UD, undetectable.

**SFigure 3.**
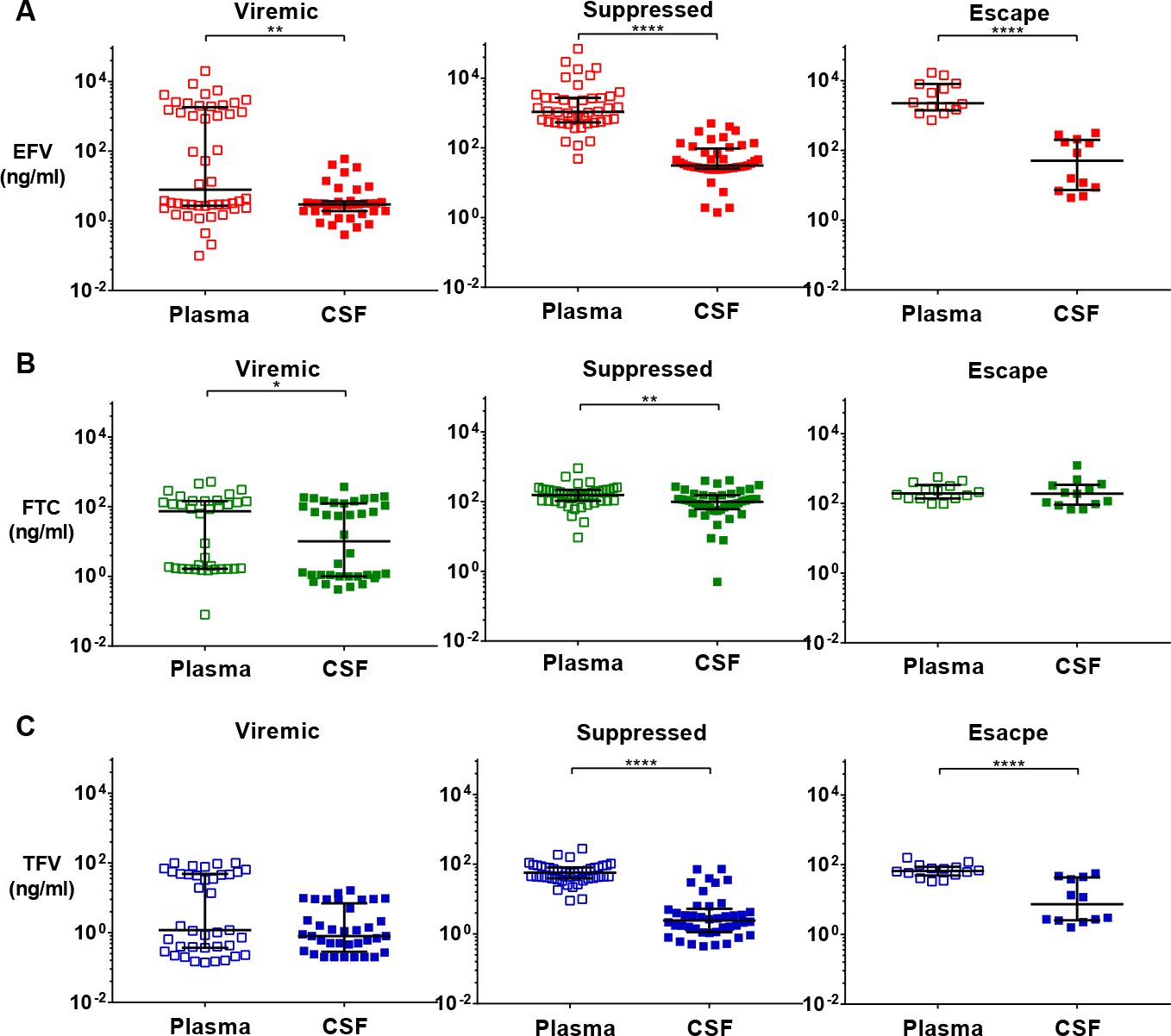
Precipitous drop in EFV levels between CSF and plasma. **(A)**, EFV concentrations in the plasma compared to CSF in viremic, suppressed and CSF Escape participants. Median EFV concentration in viremic individuals was 7.8 ng/ml (IQR 2.7-1833 ng/ml) in the plasma versus 2.9 ng/ml (IQR 1.9-3.6 ng/ml) in the CSF (*p* = 0.003). Median EFV in suppressed was 1080 ng/ml (IQR 538-2720 ng/ml) in plasma versus 31.1 ng/ml (IQR 25.2-96.3 ng/ml) in the CSF (*p* < 0.0001). Median EFV in CSF Escape was 2250 ng/ml (IQR 1403-8048 ng/ml) in plasma versus 50.7 (IQR 7.3-201 ng/ml) in the CSF (*p* < 0.0001). **(B)**, FTC concentrations in the plasma compared to CSF in viremic, suppressed and CSF Escape participants. Median FTC concentration in viremic individuals was 74.1 ng/ml (IQR 1.66-144 ng/ml) in the plasma versus 10.1 ng/ml (IQR 1.0-124 ng/ml) in the CSF (*p* = 0.04). Median FTC in suppressed was 155 ng/ml (IQR 105-218 ng/ml) in plasma versus 98.3 ng/ml (IQR 60.6-153 ng/ml) in the CSF (*p* = 0.005). Interestingly, there was no significant difference (*p* = 0.5) between median FTC in CSF Escape between plasma (194 ng/ml, (IQR 139-337 ng/ml)) and CSF (190 ng/ml, (IQR 92.1-342 ng/ml)). **(C)**, TFV concentrations in the plasma compared to CSF in viremic, suppressed and CSF Escape participants. Median TFV concentrations in viremic individuals were low with no significant difference (*p* = 0.08) between plasma (1.2 ng/ml, (IQR 0.4-48.3 ng/ml)) and CSF (0.8 ng/ml, (IQR 0.3-6.9 ng/ml)). Median TFV in suppressed was 57.2 ng/ml (IQR 39.2-81.2 ng/ml) in plasma versus 2.4 ng/ml (IQR 1.1-5.3) in the CSF (*p* < 0.0001). Median TFV in CSF Escape was 65.7 ng/ml (IQR 48.7-87.2 ng/ml) in plasma versus 7.4 ng/ml (IQR 2.6-43.1) in the CSF (*p* < 0.0001). All statistical comparisons were performed using the Mann-Whitney non-parametric test.

**STable 1:**
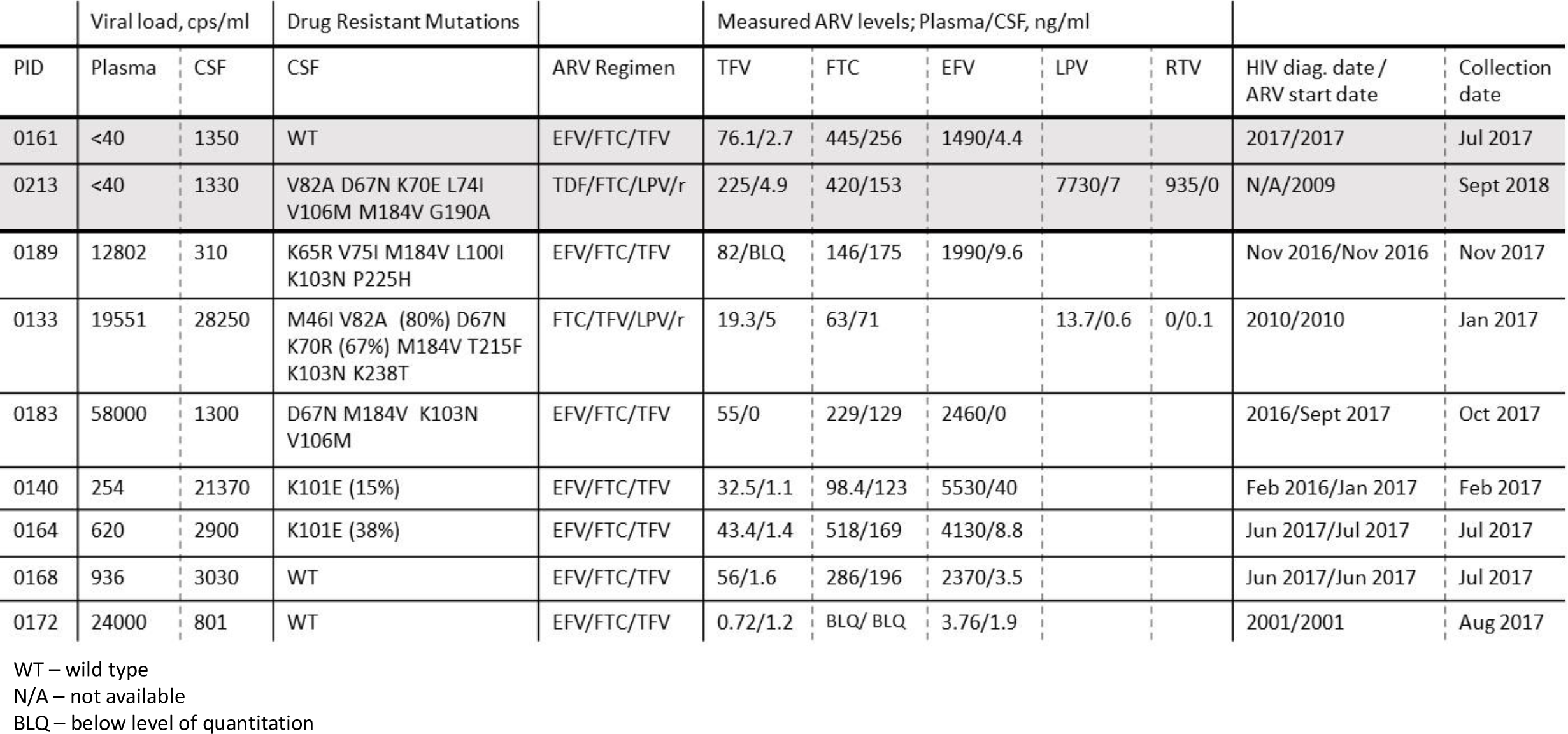
Drug Resistant Mutations in CSF from escape and viremic participants

